# WAVE1 and WAVE2 facilitate human papillomavirus-driven actin polymerization during cellular entry

**DOI:** 10.1101/2024.10.28.620484

**Authors:** DJ Fernandez, Stephanie Cheng, Ruben Prins, Sarah F Hamm-Alvarez, W Martin Kast

**Affiliations:** Department of Molecular Microbiology & Immunology and Norris Comprehensive Cancer Center, University of Southern California, Los Angeles, CA, United States; Department of Ophthalmology, Roski Eye Institute, Keck School of Medicine, University of Southern California, Los Angeles, CA, United States; Department of Pharmacology and Pharmaceutical Sciences, School of Pharmacy, University of Southern California, Los Angeles, CA, United States

## Abstract

Human Papillomavirus Type 16 (HPV16) is an etiological agent of human cancers that requires endocytosis to initiate infection. HPV16 entry into epithelial cells occurs through a non-canonical endocytic pathway that is actin-driven, but it is not well understood how HPV16-cell surface interactions trigger actin reorganization in a way that facilitates entry. This study provides evidence that Wiskott-Aldrich syndrome protein family verprolin-homologous proteins 1 and 2 (WAVE1 and WAVE2) are molecular mediators of the actin polymerization that facilitates HPV endocytosis and intracellular trafficking. We demonstrate through post-transcriptional gene silencing and genome editing that WAVE1 and WAVE2 are critical for efficient HPV16 infection, and that restoration of each in knockout cells rescues HPV16 infection. Cells lacking WAVE1, WAVE2, or both, internalize HPV16 at a significantly reduced rate. Analysis of fluorescently labeled cells exposed to HPV16 and acquired by confocal fluorescence microscopy revealed that HPV16, WAVE1, WAVE2, and actin are all colocalized at the cellular dorsal surface. We also found that HPV16 stimulates WAVE1 and WAVE2-mediated cellular dorsal surface filopodia formation during the viral endocytic process. Taken together, this study provides evidence that the HPV endocytic process needed for infection is controlled by actin reorganization into filopodial protrusions and that this process is mediated by WAVE1 and WAVE2.

**Author Summary:** Human Papillomavirus (HPV) is the most common sexually transmitted infection in the United States. While its mode of entry into cells has yet to be fully described, extensive studies indicate HPV entry occurs via a macropinocytosis-like pathway. Interestingly, more than 10 viruses enter cells via macropinocytosis-like entry, with no two viruses utilizing identical factors for entry. It is unclear whether these viruses are entering cells via the same pathway, or if the term “macropinocytosis” describes a subset of endocytic pathways. One unifying feature of entry for each of these viruses is their requirement of actin polymerization. In this study, we identify the cellular factors necessary for actin polymerization to participate in HPV endocytosis. The findings of this study are of importance to the field of virology as they may extend to the infection of other viruses. It is also of interest in cancer studies as macropinocytosis has been associated with the scavenging of nutrients and methuosis, a form of cell death in cancer cells that occurs from over- scavenging. Nanoparticle delivery can also occur via macropinocytosis. Therefore, the contribution of WAVE proteins to macropinocytosis and macropinocytosis-like endocytic events is informative to a broad audience.

## Introduction

Nearly a third of men and women worldwide are estimated to be infected by the Human Papillomavirus (HPV), a small non-enveloped DNA virus(1,2). High-risk genotypes, such as HPV type 16 (HPV16), can cause a variety of anogenital and head & neck cancers(3). While most infections are cleared by the immune response, malignancies attributed to HPV16 are the result of persistent infection, which can occur in nearly 10% of infected individuals(4).

Entry into host cells is a critical step for HPV16 infection. HPV16 has tropism for basal keratinocytes and gains access to them through micro-wounding of skin epithelia(5). Infection occurs in a complex, stepwise manner, that is initiated by binding of the virus to heparin sulfate proteoglycans (HSPGs) within the extracellular matrix (ECM). During the wound healing process, cells secrete enzymes that deconstruct HSPGs and partially cleave HPV capsids, which enables the transfer of these viral particles onto keratinocyte surfaces. Binding to keratinocyte HPV entry receptors, including but not limited to epidermal growth factor receptor (EGFR), laminin binding integrins (α6β4, α3β1), and the annexin A2 heterotetrametric protein (A2t), is enabled by the cell migration that occurs during wound healing. Consequently, intracellular signaling pathways are activated to mobilize the cytoplasmic machinery necessary for HPV endocytosis to occur(6).

The HPV16 endocytic mechanism is currently best described as incomplete. Extensive studies by our lab and others of HPV endocytosis have demonstrated that it occurs independently of clathrin, caveolin, flotillin, lipid rafts, cholesterol, and dynamin(7,8). As such, the molecules that orchestrate the inward membrane deformation observed in electron micrographs as well as the vesicle scission from the plasma membrane have yet to be identified. Biochemical studies have identified similar, but not identical, molecular mediators of macropinocytosis as contributing to HPV endocytic internalization(9). As such, HPV endocytosis has been described as “macropinocytosis-like.”(8) Macropinocytosis is characterized by the nonspecific internalization of extracellular fluid into large (0.2-5 μm in diameter) vesicles(10). Membrane protrusions that engulf cargo into macropinosomes are largely actin-driven(11). In contrast, HPV-containing vesicles are typically 0.07-0.140 μm in diameter(8). While previous studies have made clear that the presence of actin filament networks play a critical role for HPV endocytosis, the specific contribution of actin dynamics to HPV endocytosis remains understudied(8,12–15).

Actin filament participation in endocytosis has been most extensively studied within the context of clathrin-mediated endocytosis (CME)(16,17). Early studies in budding yeast observed that actin assembly components are recruited by clathrin adaptor proteins(18). Many effectors of actin assembly have been identified, including the actin-related proteins 2/3 (Arp2/3) complex, which facilitates the addition of actin monomers (G-actin) onto actin filaments (F-actin)(19,20). In CME, the Arp2/3 complex is anchored to the actin nucleation promoting factors (NPFs) Wiskott- Aldrich Syndrome Proteins (WASP and Neural-WASP), which are extensively regulated as a major mechanism that controls actin-dependent events(19). In contrast to CME, the clathrin- independent endocytic mechanism macropinocytosis, involves the direct recruitment of NPFs and the Arp2/3 complex to the transmembrane proteins that initially transmit the extracellular signal into the cytoplasm(21). However, little detail is known of the involvement of NPFs that contribute to this macropinocytosis-like endocytic mechanism.

The WASP and WASP-family verprolin-homologous (WAVE) protein family consists of nine members in mammals that have been well described in a recent review(22). These include WASP & N-WASP, WASP-family verprolin-homologous proteins 1, 2, and 3(WAVE1-3), Wiskott-Aldrich syndrome protein and SCAR homologue (WASH), WASP homolog-associated protein with actin, membranes and microtubules (WHAMM), junction-mediating and regulatory protein (JMY), and WAVE homology in membrane protrusions (WHIMP), a recently discovered family member(23). They each activate the Arp2/3 complex and couple it to G-actin through their homologous C-terminal domains(24), enabling the formation of branched actin network. The WASH functional contribution occurs downstream of the retromer complex to transport endosomes to the Golgi apparatus, and WHAMM and JMY provide actin-associated structural support to autophagosome-lysosome-Golgi apparatus vesicular trafficking(23). Very little is known of WHIMP, and WAVE3 is not expressed in epithelial tissues, so we excluded these proteins from this study. Here, we investigate the contributions of WASP and WAVE proteins to endocytosis in an HPV16 infection model. We hypothesize that the actin-driven forces generated by cell surface stimulation by HPV16 through binding to viral entry receptors occur because of specific activation of WASP- WAVE proteins. We utilize both siRNA-mediated gene silencing and CRISPR-Cas9-based genome editing to test and confirm that WAVE1 and WAVE2 are contributors to HPV endocytosis. In addition, we describe the observation of a WAVE-mediated morphological event: the formation of cellular dorsal surface actin protrusions that occur during HPV infection.

## Results

### Wiscott-Aldrich Syndrome protein family members 1 and 2 (WAVE1 and WAVE2) are critical for the infection of HPV16 in HeLa cells

Microscopy and biochemical studies have shown that HPV16 relies on actin dynamics for endocytosis and endocytic trafficking. However, the factors that respond to cell surface binding by HPV16 which guide actin polymerization intracellularly are unknown. We investigated the WASP/WAVE family of actin nucleation promoting factors for their ability to disrupt HPV16 pseudovirus infection (**Fig 1**). Out of the nine WASP/WAVE family members, evidence of cell surface activity exists for WASP, WAVE1, and WAVE2(23). The related homologue, WAVE3, also appears to function at the cell surface but this protein is not expressed in anogenital epithelial tissue(25,26). HPV16 infection has been shown to be dependent on the laminin binding integrin α6 and β4, so we knocked down the β4 subunit as a biological positive control, as this target would also prevent the post-translational processing of the α6 subunit. This proved to significantly reduce HPV infection (**S1**). The transfection process also had a minor effect on the infection rate due to cellular toxicity inherent in the transfection process. The loss of protein expression mediated by three independent siRNAs targeting either WASP, WAVE1, and WAVE2 was confirmed at the endpoint (120 h) of the infection assays (**Fig 1A, C, E, G**) by Western blotting. Here, HPV16 infection is defined experimentally by the expression of a reporter gene in cells delivered by HPV16 pseudovirions. Knockdown of WASP did not significantly affect infection in two of the three siRNAs tested as compared to cells treated with a scrambled negative control siRNA (**Fig 1B**). However, all three siRNAs targeting WAVE1 resulted in a significant reduction in infection of ∼35% (**Fig 1D**). Significant reduction in HPV16 infection of ∼40% was also seen with knockdown of WAVE2 (**Fig 1F**). Of note, western blot images depict a representative replicate.

**Fig 1.**
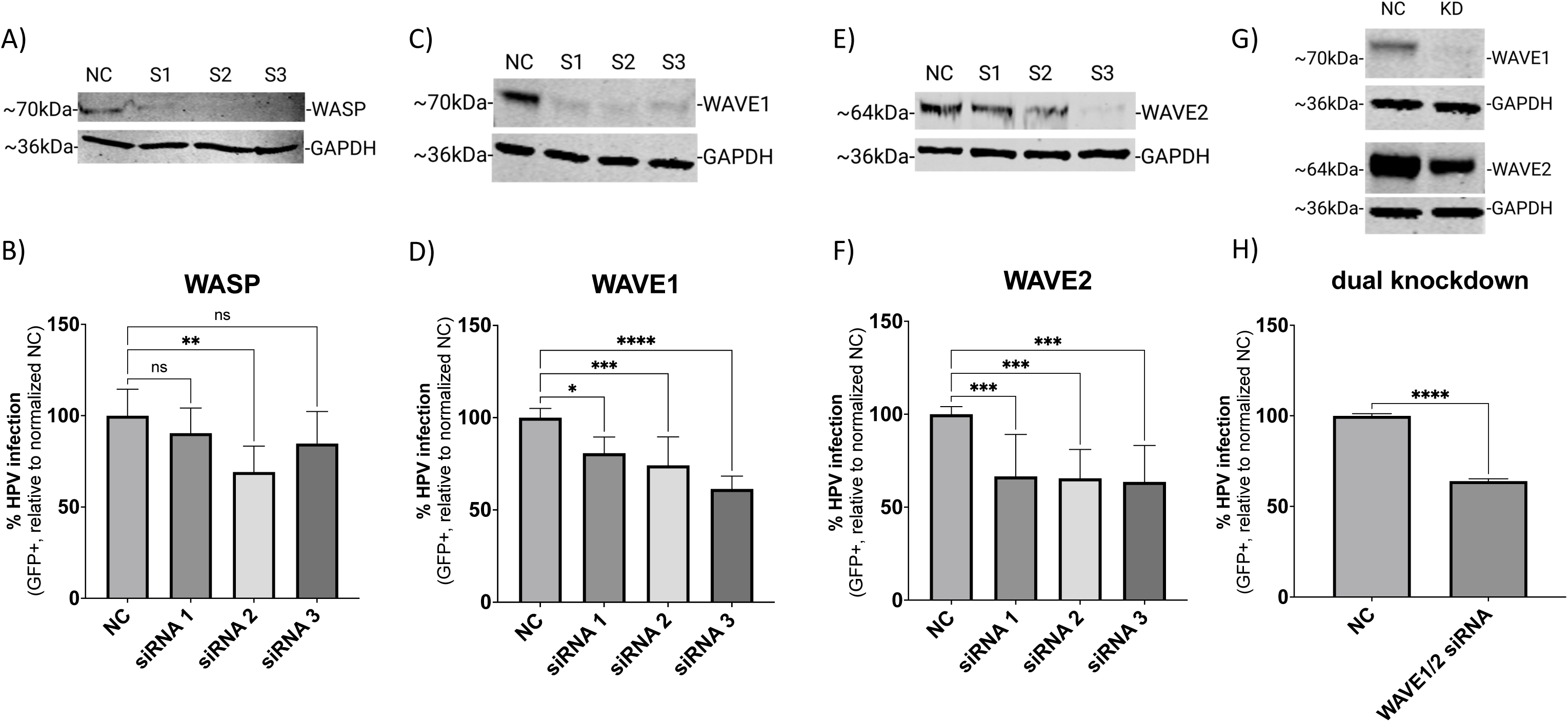
siRNA-mediated knockdown of WAVE1 and WAVE2 inhibits HPV16 infection in HeLa cells. On day 0 HeLa cells were seeded and transfected with siRNA in a 6-well microplate. On day 2, cells were collected and seeded onto a 24-well microplate to establish technical replicates. On day 3, cells were infected with HPV16 PsVs (TCID_30_) containing a GFP reporter plasmid for 48 hours. Protein expression of relevant proteins was measured via Western blotting on day 5 (A, C, E, G). NC is the negative control siRNA used in this study, while S1, S2, and S3 refer to each of three separate siRNAs used to target the indicated proteins. For panels G and H, S2 targeting WAVE1 and S3 targeting WAVE2 were employed to achieve knockdown of both proteins. Half volumes of each siRNA were combined for transfection so that the final concentration of siRNA in each experiment remained consistent. The percentage of HPV16 infected cells was also determined on day 5 (48 hours post infection) via flow cytometry (B, D, F, H). Each bar represents three biological replicates comprised of technical triplicates and show the mean %GFP+ cells ± standard deviation (n=3, normalized to WT). 1-way ANOVA with Dunnett’s multiple comparisons test was used to statistically determine significance (ns=not significant, **p<0.001, ***p<0.0001, ****p<0.0001).

According to the literature, certain morphological events such as cell migration can be facilitated by WAVE1 or WAVE2 alone, but loss of both proteins severely impairs the process(27). As such, we pooled together the S2 siRNA targeting WAVE1 and the S3 siRNA targeting WAVE2 (the final total concentration of siRNA remained 50 nM as with single knockdown experiments) as they were average performers in the infection assays. This dual knockdown yielded an infection reduction of about 40% (**Fig 1H**). While suggestive of a role for both isoforms, these results prompted an examination of whether a complete loss of function of WAVE proteins would result in a potentially more severe phenotype.

### WAVE1 and WAVE2 individually facilitate HPV16 entry

To better understand how the presence of WAVE proteins affects the HPV16 infection rate in a population of cells, we utilized CRISPR- Cas9 to generate clonal populations of cells harboring a knockout of WAVE1, WAVE2, or of both genes. Western blot analysis of WAVE protein quantification confirmed total loss of expression (**Fig 2A**). Phase-contrast imaging of cells revealed profound morphological deviations from wild- type morphology, indicating that loss of WAVE proteins affected cytoskeletal arrangement (**Fig 2B**). As compared to wild-type HeLa cells, WAVE1 knockout HeLa cells (W1KO) harbor more long, narrow projections (**Fig 2B, black arrows**). In contrast, WAVE2 knockout cells (W2KO) and the double knockouts (W1/W2KO) lack projections and instead exhibit wide lamellipodial surfaces at the cellular periphery in W2KO cells (**Fig 2B, white arrows**) or constitutively active blebbing in double knockouts (**Fig 2B, last image)**. While individual cells in WT and W1KO cell populations do not associate closely in proximity until high confluency, W2KO and W1/W2KO cells form colonies of 3-5 cells at any confluency and never establish a uniform sheet-like culture at high confluency as WT cells do. As actin polymerization is an essential function for cell survival, we tested the ability of knockout cells to proliferate normally. All knockout cell populations proliferated at an equal rate to WT cells as determined by the CyQUANT Cell Proliferation Assay (**Fig 2C**) which is particularly of note in the context of the blebbing W1/W2KO cells, which seems to be unassociated with apoptosis. Blebbing in nonapoptotic cells has been described in literature, although it is not a well-characterized phenotype(28–30). Knockout of WAVE1, WAVE2, or both proved to impair HPV16 infection in HeLa cells more severely (**Fig 2D**). While infection in W1KO cells was about 45% reduced compared to WT, knockout of WAVE2 caused a greater reduction of infection, reducing by 68% relative to WT. Importantly, there was no statistically significant difference determined between infection rate in either knockout cell population. To confirm these findings in a different cell line, we utilized B16-F1 melanoma cells lacking WAVE1, WAVE2, or both, generated by the Bruce Goode Laboratory. The resulting infection assays achieved similar results as in HeLa cells, with a significant reduction of infection in knockouts compared to WT (**Fig 2E**). However, infection in W1/W2KO B16-F1 cells was significantly lower than W1KO B16-F1 as well as W2KO B16-F1.

**Fig 2.**
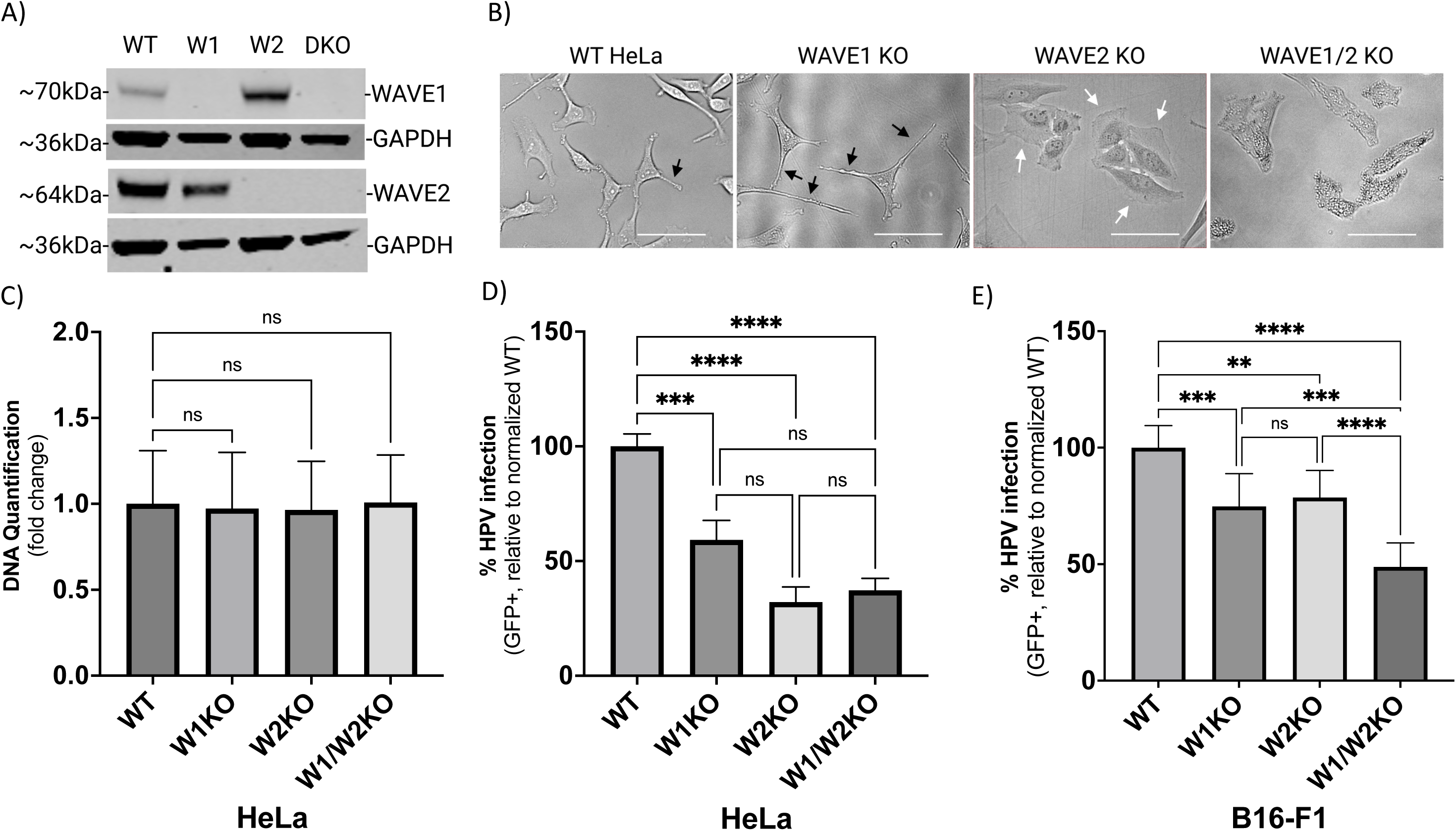
WAVE1 knockout (W1KO), W2KO, and W1/W2KO alters cellular morphology, but not proliferation, and inhibits HPV16 infection in multiple cell lines. (A) WAVE1 (W1) WAVE2 (W2) or both (DKO) proteins were knocked out in wild type (WT) HeLa cells via CRISPR/Cas9 and confirmed by Western blotting. (B) Representative phase-contrast images of WT, W1KO, W2KO, and W1/W2KO HeLa cells were taken on the FloID Cell Imaging Station (20x magnification, scale bar = 50μm). (C) W1KO, W2KO, and W1/W2KO HeLa cells were seeded in equal amounts, grown for 48 hours, and then analyzed for differences in DNA quantity via CyQUANT Cell Proliferation Assay (Thermo Fisher) compared to WT. (D and E) WT, W1KO, W2KO, and W1/W2KO HeLa or B16- F1 cells were treated with HPV16 PsVs (TCID_30_) containing a GFP reporter plasmid. The percentage of infected cells (based on GFP reporter gene expression) was measured at 48 hours post infection via flow cytometry. Background from mock infected cells was subtracted. For HeLa cells, at least 2 independent clones of each knockout were screened for consistent inhibition of HPV16 infection. Each bar represents three biological repeats comprised of technical triplicates and show DNA quantification over 48 hours (Panel C) or the mean %GFP+ cells ± standard deviation (n=3, normalized to WT) (Panels D and E). 1-way ANOVA with Dunnett’s multiple comparisons test was used to statistically determine significance (ns=not significant, **p<0.001, ***p<0.0001, ****p<0.0001).

### Restoring WAVE1 or WAVE2 protein expression in KO HeLa cells rescues HPV16 infectivity

To confirm that the observed inhibition of infection occurred due to loss of WAVE1 and/or WAVE2 protein expression, we utilized lentiviral vectors to restore WAVE1 or WAVE2 activity back to KO HeLa cells. We confirmed via Western blotting that KO cells were expressing WAVE1 (W1 R) or WAVE2 (W2 R) (**Fig 3A and C**). To assess HPV16 infection in these cells, we infected cells with HPV16 pseudovirions as previously described. The resulting infection rate in W1 R cells was increased 76% compared to WT (**Fig 3B**). Similarly, expressing WAVE2 in W2KO cells resulted in a 112% increase in infection (**Fig 3D**).

**Fig 3.**
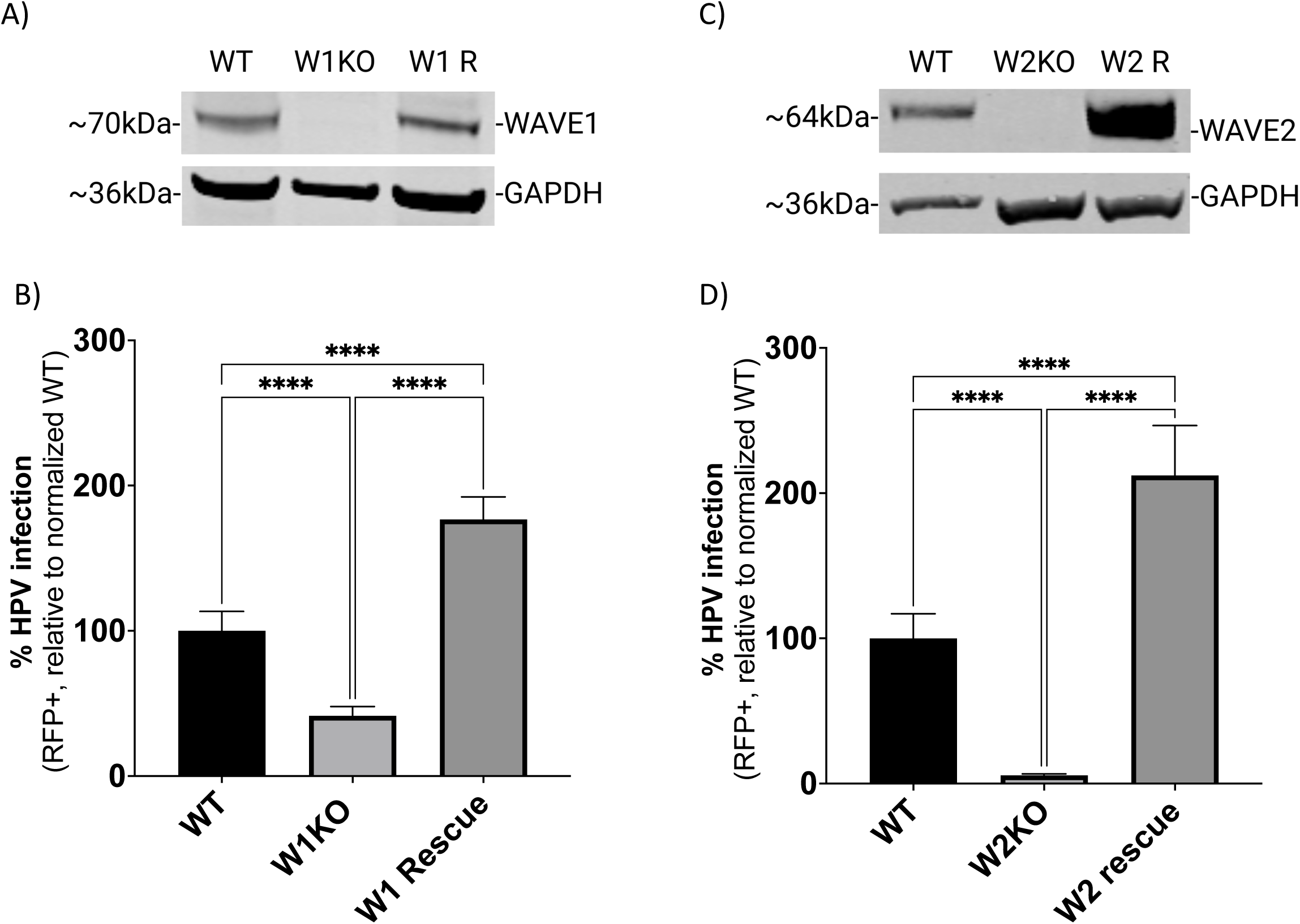
HPV infectivity is functionally recovered by WAVE1 or WAVE2 expression in HeLa cells. (A and C) WT and W1KO or W2KO cells were transduced with a mammalian gene expression lentiviral control vector or a vector containing either GFP-WAVE1 or GFP-WAVE2, respectively (Vector Builder). Transduced cells received an antibiotic resistance gene and underwent selection. (B and D) WT, KO, and cells with WAVE protein expression restored were treated with HPV16 PsVs (TCID_30_) containing an RFP reporter plasmid. The percentage of infected cells (RFP reporter gene transduction) was measured at 48 hours post infection via flow cytometry. Background from mock infected cells was subtracted. Each bar represents three biological repeats comprised of technical triplicates and show the mean %RFP+ cells ± standard deviation (n=3, normalized to WT). 1-way ANOVA with Dunnett’s multiple comparisons test was used to statistically determine significance (ns=not significant, **p<0.001, ***p<0.0001, ****p<0.0001).

### WAVE1 and WAVE2 are required for proficient HPV16 internalization and endocytic trafficking

We next investigated if the reduction in HPV infection due to the loss of WAVE1 and/or WAVE2 could be explained by a reduction in HPV cell surface binding, as cell surface receptors may require WAVE-mediated actin dynamics to establish and maintain homeostatic expression levels. Contrarily, in W1KO and W2KO cells, there was an apparent increase in the average number of particles bound to the cell surface although it was not statistically significant (**Fig 4A**). However, the elevated number of particles on W1/W2KO cells did reach significance. These binding assay results corroborated the result of our internalization assay (**Fig 4B**). To assess HPV16 internalization, WT and KO cells were infected with HPV16 virus-like particles (VLPs) conjugated with pHrodo, a pH-dependent rhodamine dye that increases in fluorescence accordingly with decrease in pH, as occurs during endocytic trafficking through increasingly low pH membrane compartments, which we and others have shown previously(31–33). HPV16 endocytosis is known to follow a retrograde endosomal trafficking pattern and travel through the Golgi apparatus and endoplasmic reticulum before reaching the nuclear compartment, a process that has been described to occur over 7-8 hours (34). We found that the rate of increase in signal intensity was significantly slowed over the 7 h time course in W2KO and W1/W2KO, but the observed reduction in W1KO cells was not significant (**Fig 4B**). We next investigated if the decreased rate of signal intensity was due to an inability of particles to travel along the retrograde endosomal pathway. We approached this by utilizing confocal microscopy in a time course imaging assay. Cells were infected with PsVs and, over the course of 8 h, examined to evaluate the colocalization between HPV16 and organelles involved in HPV retrograde transport. Although the abundance of HPV16 was decreased in internal compartments, the time course of colocalization of the virus with membrane compartments including early endosomes (EEA1), multivesicular bodies (VPS25), Golgi apparatus (Golgin97), and endoplasmic reticulum (SERCA2) and were unaffected with one exception (**S2**). In W2KO cells as early as 2 hours of infection, we detected increased HPV16 colocalization with a lysosomal marker (LAMP1) in all knockout cells as compared to wild type cells (**Fig 4C**). Taken together, there was an increase in the number of HPV16 particles bound to cells lacking both WAVE1 and WAVE2, while those cells also internalized viral particles more slowly and trafficked an increased number of them towards lysosomes.

**Fig 4.**
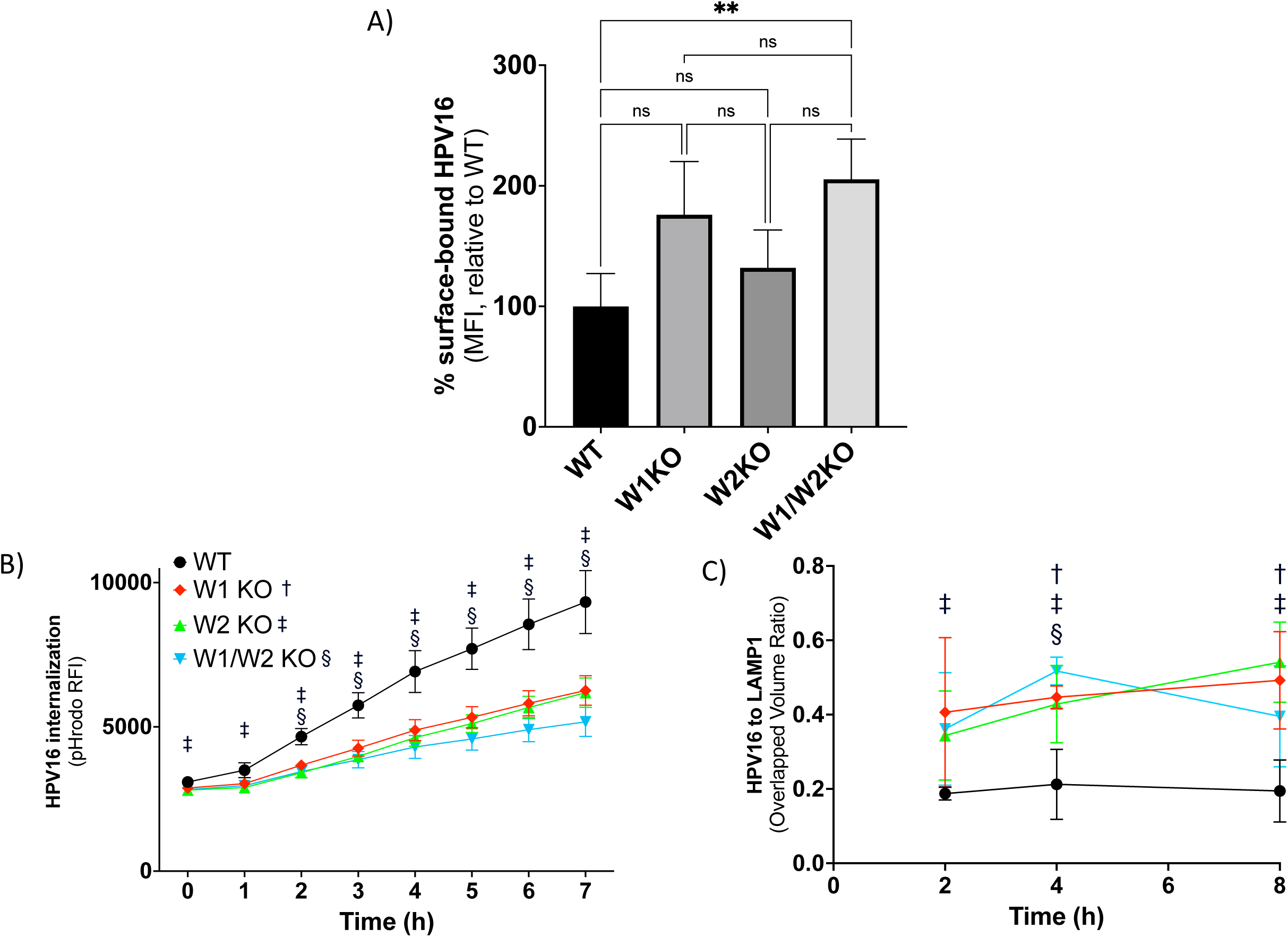
W1KO, W2KO, and W1/W2KO increase HPV16 surface binding, reduce rate of internalization, and increase trafficking of particles to the lysosome. (A) To assess the ability of HPV16 to bind its coreceptors, WT or KO cells were cooled to 4°C for 0.5 h to inhibit endocytosis. Cells were then transferred to ice and saturated with HPV16 VLPs (10 μg/1E6 cells) in serum-free media for 1 hour at 4°C. Cells were collected via scraping over ice and then subjected to immunostaining. The quantity of surface-bound HPV16 was analyzed by flow cytometry. Results show the mean fluorescent intensity (MFI) ± standard deviation, normalized to WT. (B) Cells were treated with pHrodo-labelled HPV16 VLPs (5 μg/1E6 cells) for 7 hours at 37°C and measured each hour via plate reader (BMG Labtech). Results show the mean MFI ± standard deviation. (A) and (B) represent three biological repeats comprised of technical triplicates. (C) Cells were cooled to 4°C for 0.5 h prior to the addition of HPV16 VLPs (0.5 μg/1E6 cells) diluted in ice-cold media and incubated together at 4°C for 1h. Next, cells were transferred to 37°C for either 2, 4, or 8h and subsequently fixed with 4% paraformaldehyde. Sample next underwent immunostaining for LAMP1 and HPV16, with a nuclear counterstain (DAPI). At least 5 Z-stacks were imaged via confocal microscopy from each of 3 biological repeats (∼15 Z-stacks total per sample type with a minimum of 15 cells per condition). The quantification of the extent of colocalization between HPV16 and LAMP1 was measured by determining the overlapped volume ratio of voxels using Imaris. Results are depicted as the mean overlapped volume ratio ± standard deviation. Statistics: (A) 1-way ANOVA with Dunnett’s multiple comparisons test was used to statistically determine significance (ns=not significant, **p<0.001). (B & C) Multiple unpaired t-tests were conducted using the Holm-Šídák method for each time point between WT and KOs. †, ‡, §, symbols correspond with W1KO, W2KO, and W1/W2KO, respectively, and indicate p<0.05.

### HPV16 colocalizes with WAVE1 and WAVE2 at the cellular dorsal surface

Virus internalization experiments in **Fig 4B** suggested that the functional contribution from WAVE1 and WAVE2 toward HPV16 entry began within the first 2 h of virus addition, since inhibition of virus uptake was significant after 2 h in cells lacking either or both proteins. Accordingly, we utilized an imaging approach to investigate if WAVE1 and WAVE2 colocalized with HPV16 within the endocytic timeframe. To do so, we employed WT HeLa cells expressing GFP-actin and cooled them from 37°C to 4°C for 0.5 h to inhibit endocytosis. We then added HPV16 PsVs to cells for 1 hour at 4°C to facilitate surface attachment. Cells were then returned to 37°C for 30 minutes prior to fixation, to allow time for cellular processes such as endocytosis to re-initiate. Samples were then fixed and immunolabeled to visualize HPV16 along with WAVE1 **(Fig 5A)** or WAVE2 **(Fig 5C)**, and actin. Z-stacks were imaged to identify the cellular dorsal surface. We found that WAVE1 colocalized mostly with cortical actin and was less present in lamella **(Fig 5A image 3)**, while WAVE2 colocalized with both cortical actin as well as at the leading edges of lamellipodia **(Fig 5C image 13)**. We observed HPV16 particles bound particularly at the cellular dorsal surface, above and surrounding the area of the nuclear stain, and fewer particles at the cellular periphery **(Fig 5 images 4 & 14)**. We also found that at locations on the dorsal surface that harbored HPV16, there appeared to be an enrichment of fluorescence intensity signal from the actin GFP-tag as well as from the immunostained WAVE2 **(Fig 5 images 2 & 12, and 13 respectively).** To assess the spatial relationship between WAVE proteins, actin, and HPV16 particles, we generated images that depict colocalized voxels between signals **(Fig 5B & D)**: WAVE proteins and actin **(Fig 5 images 6 & 16)**, HPV16 and actin **(Fig 5 images 7 & 17)**, HPV16 and WAVE proteins **(Fig 5 images 8 & 18)**. We also overlaid the images depicting the colocalization of HPV16 and actin with the images of the colocalization between HPV16 and WAVE proteins **(Fig 5 images 9 & 19)** to appreciate the clear distinction of points in which HPV16, WAVE proteins, and actin are all colocalized, which appear directly above and surrounding the nucleus, and are most clearly represented in **Fig 5 images 10 and 20**.

**Fig 5.**
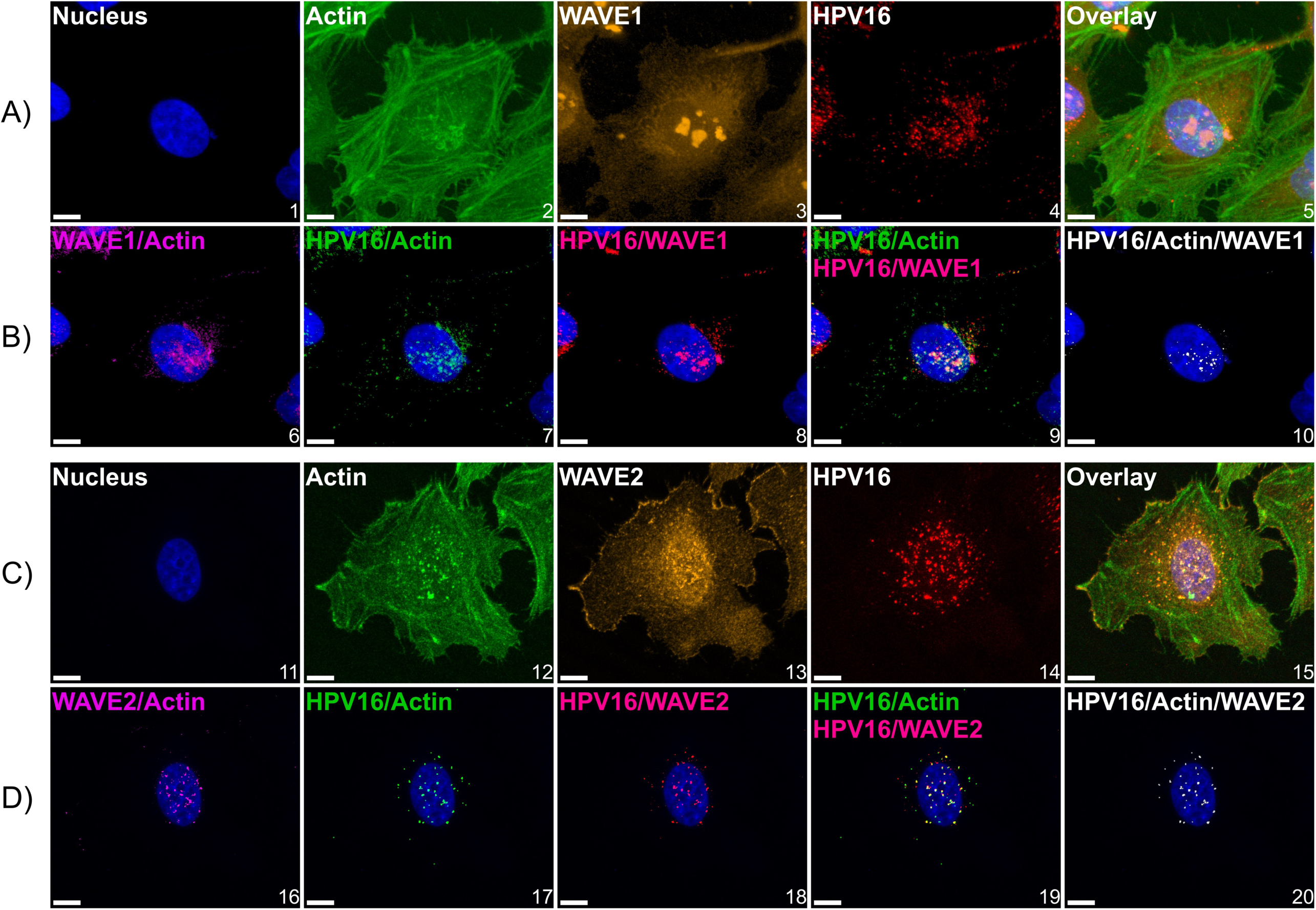
HPV16 colocalizes with actin and WAVE proteins at the cellular dorsal surface. WT HeLa cells expressing LifeAct-GFP seeded in chambered microscope slides were first cooled from 37°C to 4°C for 0.5 h to inhibit endocytosis prior to the addition of HPV16 VLPs (10 ng/1E6 cells) in ice cold media for 1 hour. Cells were then returned to 37°C for 10 minutes prior to fixation with 4% paraformaldehyde for 10 minutes at room temperature, which was the temperature for subsequent steps. Samples were then permeabilized with 0.1% Triton X-100, blocked with 1% BSA, and immunostained against HPV16 L1 and (A) WAVE1 or (C) WAVE2. Hoescht 33342 was added during secondary antibody addition as a counterstain. Z-stacked images were generated via laser scanning confocal microscopy. (A and C) maximum intensity projections of Z-stacks of images depicting candidate cells. The color channels are labeled at the upper left of each image. (B and D) to analyze the spatial relationship between signals, we utilized Imaris 10.1.1 Microscopy Image Analysis Software (Oxford Instruments). Briefly, a “surface” was created for each signal, which is an Imaris segmentation algorithm. Surfaces were generated to provide object-object statistics. Parameters included the smoothing of surface details to 0.2 um with the method of absolute intensity thresholding. Background signal was subtracted through voxel size filtration (voxels smaller than 10 were excluded). Next, colocalization between channels was determined by the colocalization tool. Colocalized voxels (as determined by a Manders’ coefficient of 1) between surfaces were determined by first thresholding images to include true signals and restrict noise. New channels were then created of colocalization voxels. For both conditions, 3 fields containing 5-15 cells across 3 biological replicates were imaged. Scale = 10 µm.

### HPV16 stimulates dorsal surface membrane protrusions

Previous studies have shown that HPV16 induces peripheral filopodia and utilizes them for retrograde transport towards the cell body prior to endocytosis(14,15). Additionally, WAVE proteins have been implicated in generated dorsal surface protrusions(35). As we found that HPV16 colocalized with actin, WAVE1, and WAVE2 at the dorsal surface, we investigated if PsVs also stimulated actin protrusions there. To approach this, cells were treated with CellLight Actin-GFP, Bacmam 2.0 upon seeding into 8-well chamber slides 24 h prior to HPV16 stimulation for 0.5-2 h. Cells were then immediately fixed, and z-stacks of images were taken via confocal fluorescence microscopy, stitched to produce a volume view, and rotated to a perspective view to appreciate the actin protrusions in the Z- direction **(Fig 6A, 0.5 hour infection depicted)**. In the absence of HPV16, HeLa cells expressed ≤ 1 dorsal surface or peripheral filopodia. However, HPV16-stimulated WT cells expressed dorsal surface membrane rufles after 30 minutes **(Fig 6A image 8)**. We did not find published methods to analyze actin protrusions in the Z-direction, so those that we observed were identified using the following criteria: they had to extend from within and above the nuclear perimeter, and they had to extend greater than 1 μm above the nuclear stain. The microscopy image analysis software, Imaris, was used to measure the length of protrusions in the Z-direction using the measurement tool. We have included an example of a cell with measured protrusions **(S3)**. These actin protrusions varied in length, with the longest found to be 6 µm **(Fig 6B)**. On average, protrusions were between 1.5-2 µm long.

**Fig 6.**
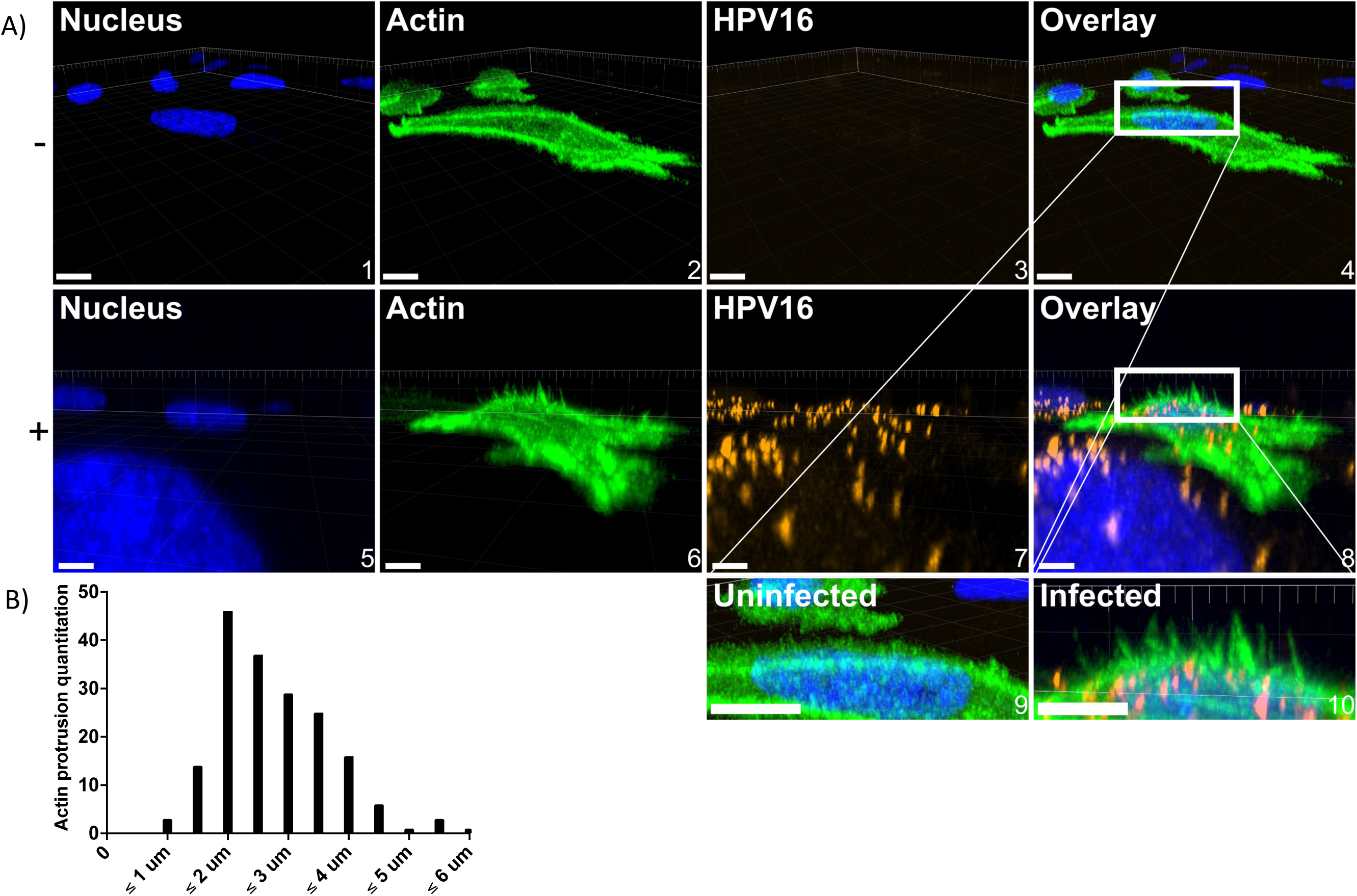
WT HeLa cells stimulated by HPV16 express dorsal surface actin protrusions. Cells were prepared as described in Fig 5; however, cells were not permeabilized during immunostaining. A) either untreated (top) or HPV16 infected HeLa cells (10 ng/1E6 cells) (bottom) treated with CellLight Actin-GFP were imaged via laser scanning confocal microscopy to obtain Z-stacks. Z-stacks were then stitched together and rotated to view the XZ oriented volume. Overlaid images (4 and 8) include a white box to indicate where dorsal surface actin protrusions appear. Images 9 and 10 depict what is in the white boxes but scaled up. Scale = images 1-4, 10 µm; images 5-8, 6 µm. 20 cells were analyzed per condition. B) Actin protrusion quantification was done using Imaris. The draw tool was utilized within the Surpass Tree Item Volume with the FITC channel selected. Spheres (points) were added at the base of actin protrusions, which stemmed perpendicularly from the actin cortex. The base of filopodia was determined to be the vertex of where the filopodia and the actin cortex meet. Next, a sphere (point) was added to the distal end of the filopodia as determined by fluorescence intensity. The distance between spheres was then determined.

### WAVE1 and WAVE2 are necessary for HPV16-driven dorsal surface actin protrusions

Since HPV16 stimulated dorsal surface actin protrusions in WT cells, we repeated the above imaging assay in W1KO, W2KO, and W1/W2KO cells to investigate differences in HPV16-induced dorsal surface protrusions compared to WT (**Fig 7**). Again, we found that almost all observed untreated WT cells had smooth dorsal surfaces, while HPV16 stimulated WT cells expressed dorsal surface protrusions **(Fig 7, images 1 & 5, respectively)**. However, both treated and untreated knockout cells had smooth surfaces. Indeed, knockout cells expressed few to zero dorsal surface protrusions that were quantifiable **(Fig 8)**. Collectively, this data indicates that HPV stimulates WAVE-mediated dorsal surface actin protrusions.

**Fig 7.**
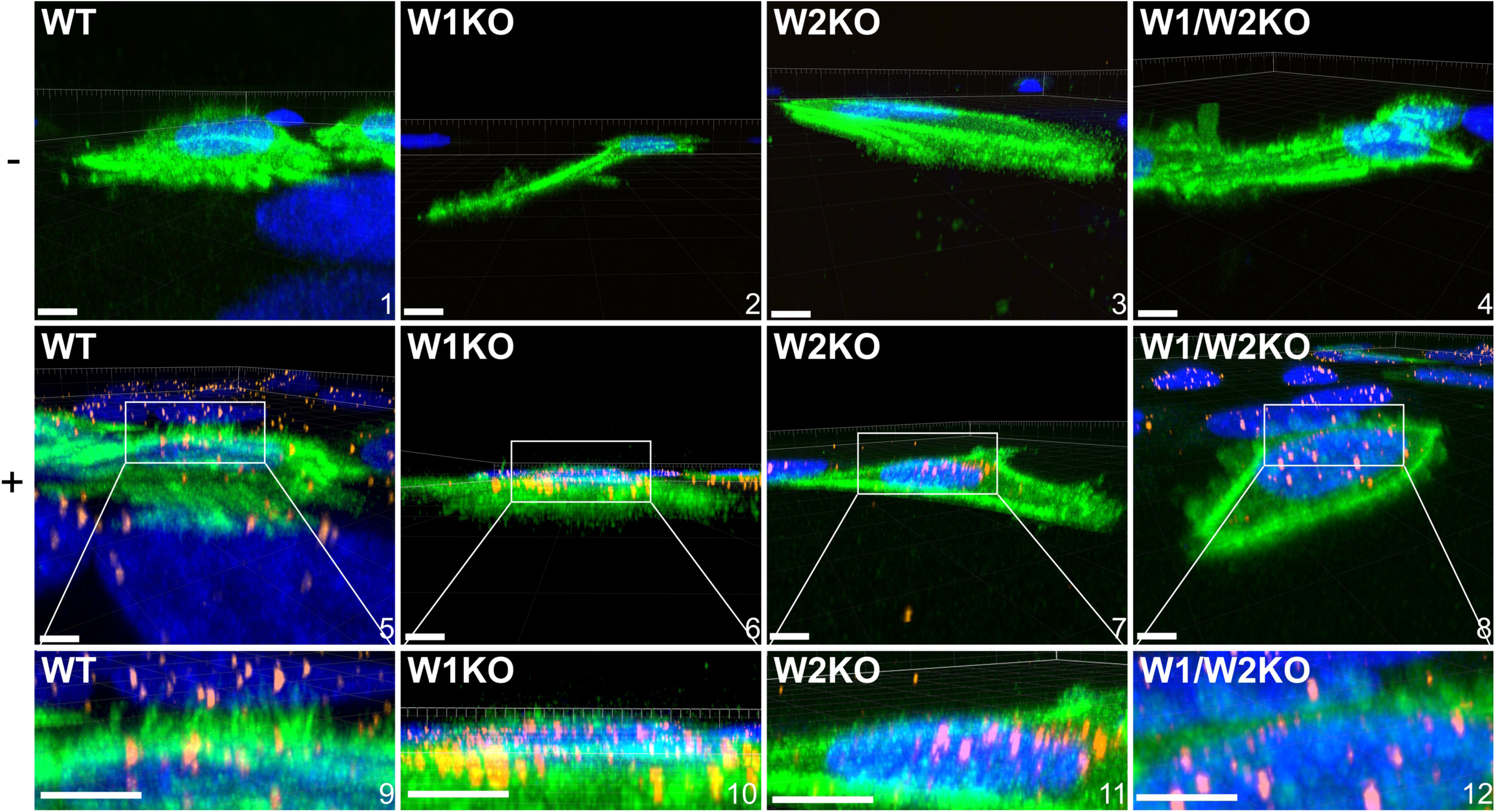
Knockout of WAVE1, WAVE2, or both, prevents HPV16 stimulated HeLa cells from expressing dorsal surface actin protrusions. Cells were prepared as described in Fig 5 however, cells were not permeabilized during immunostaining. Either untreated (top row, - symbol) or HPV16 infected WT, W1KO, or W1/W2KO HeLa cells (10 ng/1E6 cells) (middle row, + symbol) treated with CellLight Actin-GFP were imaged via laser scanning confocal microscopy to obtain Z-stacks. Z-stacks were then stitched together and rotated to view the XZ oriented volume. Scale: images 1, 2, 4-7 = 10 µm; image 3 = 8 µm; image 8 = 14 µm. 22 cells were analyzed per condition.

**Fig 8.**
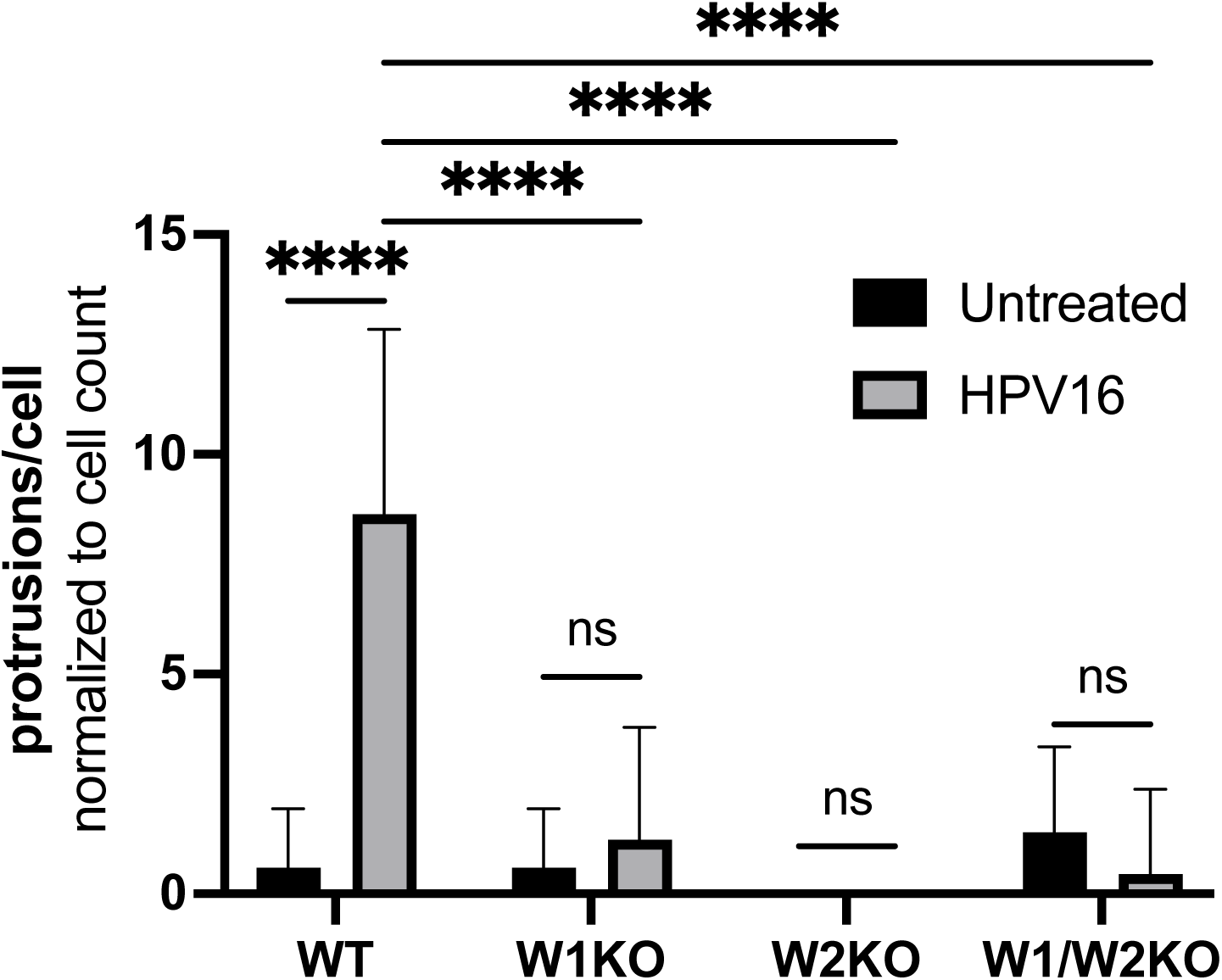
Knockout of WAVE1, WAVE2, or both, results in a significant reduction of dorsal surface actin protrusions. Dorsal surface actin protrusions were quantified using the same method as described in Fig 6. The graph depicts the average number of protrusions per cell ± standard deviation. Statistics: 2-way ANOVA with comparison of means was used to statistically determine significance, corrected for multiple comparisons using Tukey’s test (ns = not significant, ****p<0.0001).

## Discussion

The elusiveness of HPV endocytosis is exemplified by two conflicting observations: 1) that various biochemical analyses show that aspects of internalization resemble macropinocytosis; but 2) the size of typically observed nascent HPV-containing endosomes are smaller than macropinosomes(36). As such, it is understood from the literature that HPV entry occurs via a novel endocytic mechanism(6). Other viruses that utilize macropinocytosis reveal wide heterogeneity in the components necessary for endocytosis to occur(37). One unifying feature for each of these viruses, including HPV, is that their endocytosis requires actin; however, data in the literature is mostly limited to studies of the inhibition of actin polymerization or depolymerization during viral entry, without specific insight into the molecular mechanism(s) controlling the localized response of actin-driven force to the stimulation of HPV(8). We thus, sought to examine the role of actin polymerization and identify the factors that facilitate HPV entry into cells.

Our results indicate that WAVE1 and WAVE2 are the actin nucleation promoting factors that HPV16 triggers to facilitate endocytosis and subsequent infection in HeLa cells, which are the most common cell type used in HPV entry and trafficking studies. As WASP and N-WASP are recruited to sites of clathrin-mediated endocytosis but HPV endocytosis is known to be clathrin-independent, our finding that knockdown of WASP is inconsequential to HPV infection was expected (**Fig 1B**)(8). As a biological positive control, we targeted integrin β4, a known entry receptor for HPV16, for siRNA-mediated knockdown, yielding a reduced infection (S1)(38). HPV16 utilizes several additional entry receptors, including EGFR, A2t, and CD63(39). The reduced infection rate attributed to KD of one protein’s expression among a group that facilitates HPV16 entry thus set an important benchmark for our studies and led us to explore what might result from perturbing other single targets. Despite the incomplete knockdown of WAVE1 and WAVE2 with siRNAs, HPV16 infection was blocked significantly (**Fig 1**). Previous studies have struggled to experimentally distinguish WAVE1 from WAVE2, and we also encountered this challenge(27,40). To try and mitigate this, we decided to ablate their functions individually and together.

To better examine the functional consequence of deleting WAVE proteins on HPV16 infection, we used CRISPR-Cas9 to generate HeLa cells completely lacking WAVE protein expression individually and together (**Fig 2**). The resulting cells featured important morphological deviations from WT that are related to macropinocytosis (**Fig 2B**)(9,41). WAVE2 loss-of-function revealed changes in lamellipodia, which are structures that can support endocytosis, particularly during cell migration(42,43). Absence of WAVE2 resulted in lamellipodia evident on two sides of the cells with more tapered ends, or lamellipodia fully surrounding cells. Additionally, W1/W2KO cells constitutively expressed blebs which are induced by some viruses that activate macropinocytosis(44,45). It is of note that these blebs occur in healthy W1/W2KO HeLa cells which are not apoptotic, as evidenced by their normal proliferation rate.

While our study is the first to positively associate WAVE1 and WAVE2 with HPV entry, it is not the first to address their role. During our research, a preprint found that siRNA-mediated knockdown of WAVE1 and WAVE2 did not affect HPV infection(46). However, we found that both knockdown and CRISPR-Cas9-mediated knockout of WAVE1 and WAVE2 resulted in significant reduction to the rate of HPV16 infection in multiple infection models. The conflicting results between our positive siRNA results and the negative results depicted in the supplementary material in the preprint(46) is most likely attributed to the inherent variability in the process of siRNA transfection (which is evident in **Fig 1** of this manuscript) followed by our differing infection model designs. While this data is not shown here, we also confirmed that knockdown of WASH results in a significant reduction of HPV16 PsV infection in HeLa cells(46). For this study, we focused on the WASP-WAVE family proteins that have been established in the literature to function at the cellular surface.

In addition to generating our own KO cells, we obtained a second knockout cell line in B16- F1 cells to test infection in multiple cell lines. We anticipated a possible synergistic effect on infection in the double knockout cells. Indeed, in B16-F1 cells, W1/W2KO were significantly less infectable than the single knockouts (**Fig 2E**). However, in HeLa cells, WAVE2 seemed to contribute more to the blockage of infection (**Fig 2D**). This apparent bias in the effects of a loss of WAVE2 could be related to the fact that it is 9x more abundant in epithelial tissue than WAVE1(27). The different results between B16-F1 melanoma cells and HeLa cervical adenocarcinoma cells could be due to cell type specific differences in protein abundance(47). It is still unclear whether WAVE1 or WAVE2 function in tandem, redundantly, and/or if they control distinct cytoskeletal events. Importantly, the impairment of infection was prevented by the re-expression of each WAVE protein back in the KO cells (**Fig 3**). Surprisingly the re-expression resulted in higher rates of infection compared to WT in both W1KO and W2KO cells.

Cells lacking WAVE1, WAVE2, or both, show a trend towards an increased number of HPV particles bound to the cell surface when endocytosis is impaired, an effect which is significant for the double KO (**Fig 4A**). This effect may be due to altered internalization and recycling of HPV surface receptors caused by impaired actin dynamics, which increase one or more of the surface receptors. The elevated number of particles bound to the cell surface also may accumulate due to impaired HPV endocytosis. This is suggested by our internalization data which shows that particles move through the endocytic pathway more slowly in knockout cells (**Fig 4B**). Our colocalization study suggested that the HPV16 largely traffics similarly through organelles during endocytic trafficking in the absence of WAVE isoforms, suggesting that movement of the HPV, once internalized, occurs independently of WAVE proteins (**S2**). We noted an increase of HPV localized to the lysosome associated with loss of either WAVE isoform (**Fig 4C**).

After establishing that WAVE proteins contribute to the infectivity of HPV16 and that they are required for proficient endocytic trafficking, we investigated whether WAVE proteins could be recruited by HPV16 once it was bound to the surface (**Fig 5**). Interestingly, the HPV bound at the cell periphery was only weakly colocalized with actin, and even less so with either WAVE isoform. At the cellular dorsal surface, however, HPV16 was colocalized with both actin and WAVE proteins. The relevance of this observation lies in our understanding of HPV’s macropinocytosis-like endocytic process and the cell & molecular biology of macropinocytosis. While there exist only limited studies on the actual endocytic event, it is understood that HPV entry can occur at the cell periphery or at the cellular dorsal surface(36,48). Once bound to cells, most HPV particles have been observed to traffic towards the cellular dorsal surface and undergo asynchronous endocytosis within the first two hours of infection(49). Macropinocytosis can occur either at the leading edge of lamellipodia (cellular periphery), or at the dorsal surface, and while these differences are not well understood, there is evidence that cells undergoing migration use macropinocytosis at the dorsal surface to recycle adhesion molecules and other surface receptors en masse(50). HPV entry occurs *in vivo* in migrating basal keratinocytes(39). Macropinocytosis, which features actin-based membrane rufling, is critical for cellular migration to occur(11). WAVE proteins, specifically WAVE1, have been described to mediate membrane ruffling, which can occur at the periphery or at the dorsal surface, and while there is less data implicating WAVE2 in membrane rufling, the coupling of WAVE1 and WAVE2 in cellular events suggests that they likely both contribute to this process(40). In summation of our data and the literature, our observation of HPV16 particles colocalizing with actin and WAVE proteins at the dorsal surface at timepoints relevant for HPV endocytosis implies that WAVE1 and WAVE2 are involved in the endocytic event. This model was further supported by our observation of dorsal surface filopodia stimulated by HPV16 (**Fig 6**). A recent article has described the observation that HPV stimulates peripheral filopodia(14). To investigate the dorsal surface for a similar effect of HPV stimulation, we generated Z-stack images covering the full height of the cells and stitched them to observe their full volume in the XZ and YZ orientation. We found that there were indeed dorsal surface actin protrusions clustered directly over the nucleus in WT cells exposed to HPV while unstimulated WT cells lacked protrusions almost entirely. Importantly all knockout cells lacked protrusions as well, implying that the HPV-stimulated actin protrusions at the dorsal surface are WAVE1 and WAVE2 mediated (**Figs 7 and 8**). This data suggests that events occurring at the cellular cortex involve spatial specificity(51).

We conclude that the formation of dorsal surface actin protrusions is related to HPV endocytosis and is a prerequisite for the infectious entry of HPV. Importantly, one study investigating entry of HPV16, 18, and 31 found their endocytic entry sites at the identical subcellular location where we observe HPV16, actin, and WAVE proteins(48). It is likely that at least HPV genotypes 18 and 31 also require WAVE proteins for infection. In summary, our data indicate that cells lacking WAVE proteins do not form dorsal surface actin protrusions, accumulate particles on the surface, slowly internalize particles in what could be an alternative pathway, and partially shuttle particles to the lysosome where they are degraded.

WASP-WAVE proteins are becoming increasingly recognized for their roles in infection. *Shigella flexneri, Chlamydia trachomatis, and Escherichia coli* have been demonstrated to recruit N-WASP to facilitate entry and actin-based motility within a cell(52–54). Evidence suggests that the parasite *Trypanosoma cruzi* recruits N-WASP and WAVE2 during entry(55). There is also evidence for the involvement of WASP in the infection of vaccinia virus, and WASP and WAVE2 contribute to HIV-1 infection(56,57).

We propose as a model that HPV stimulation of its cell surface receptors recruits and activates WAVE1 and WAVE2 via MAPK and PI3K signaling, which have been proven to activate both HPV and WAVE proteins(58–60) and that their activation results in dorsal surface actin protrusions that are necessary for HPV endocytosis. In conclusion, this study provides the first evidence for the involvement of WAVE proteins in the endocytosis of HPV.

## Materials and Methods

### Cell Culture

HeLa cells (CCL-2, ATCC) isolated from cervical adenocarcinoma and derived knockout cell lines generated in this study were maintained in Iscove’s Modified Dulbecco’s Medium (Gibco) supplemented with fetal bovine serum (10%; Omega Scientific), 2- mercaptoethanol (0.05mM; Gibco), and gentamycin (50 units/ml; Gibco) at 37°C with 5% CO_2_ and 95% relative humidity. Two clones for each single and double knockout condition were screened in infection assays and we confirmed similar phenotypes between clones. As such, a single clone reflecting each knockout is described in this study. B16-F1 cells (CRL-6323; ATCC) and derived knockout cell lines generated by CRISPR/Cas9 were kind gifts from Dr. Bruce Goode (Brandeis University) and were cultured in DMEM (4.5 g/l glucose; Gibco) supplemented with L-glutamine (2 mM; Gibco), fetal calf serum (10%), gentamycin (50 units/ml) and HEPES (10 mM; Gibco).

### CRISPR/Cas9 gene editing

TrueCut Cas9 Protein V2 (ThermoFisher Scientific) was utilized to introduce CRISPR/Cas9-mediated frameshift indels. The following predesigned synthetic sgRNA sequences were used to target WAVE1 and WAVE2, respectively: 5’-TCTTGCGATCGAAAAGCTGC- 3’ and 5’-TGAGAGGGTCGACCGACTAC-3’. TrueGuide sgRNA HPRT1 was used as a positive control. Cas9 and sgRNA were combined with CRISPRMAX (ThermoFisher Scientific) for transfection and incubated for 48 h. Monoclonal cell populations were generated through limited dilution and subsequently underwent Sanger Sequencing to verify gene disruption. Protein expression was analyzed via Western blotting.

### Clonal proliferation analysis

The doubling rate of knockout clonal populations were determined via trypan blue exclusion as well as the CyQUANT Cell Proliferation Assay Kit (Invitrogen). WT, WAVE1 KO, WAVE2 KO, and WAVE1/WAVE2 KO cells were grown for 48 h, collected with Trypsin-EDTA, diluted 1:1 with trypan blue stain (Invitrogen), and viable cells were counted. The CyQUANT Cell Proliferation assay was used according to the manufacturer’s protocol. Fluorescence of dye-bound DNA was measured using the Clariostar plate reader (BMG Labtech). DNA was quantified by comparison to a DNA standard curve.

### Protein overexpression

WAVE1 and WAVE2 rescue clones were generated from WAVE1 KO HeLa cells and WAVE2 KO HeLa cells transduced with lentivirus containing eGFP-WAVE1 and eGFP- WAVE2, respectively and using puromycin selection (Vector Builder, Chicago, IL). Cells underwent transfection for 48 h before puromycin was added. After 7 days, monoclonal populations were generated using a dilution series. WAVE1 and WAVE 2 levels were quantified via Western blotting and expression of eGFP-WAVE1 and eGFP-WAVE2 fluorescence was determined using flow cytometry. The selected clones proliferated at a comparable rate to WT, WAVE1 KO and WAVE2 KO HeLa cells.

### Western blotting

Cell lysates were prepared by utilizing Pierce IP lysis buffer supplemented with HALT protease inhibitor cocktail according to manufacturer’s protocols (ThermoFisher Scientific). Samples of lysates were mixed with NuPAGE LDS Sample Buffer and Reducing Agent (ThermoFisher Scientific) and boiled for 10 minutes before being added to NuPAGE 10% Bis-Tris Mini Protein Gels immersed in NuPAGE MOPS SDS Running Buffer supplemented with NuPAGE Antioxidant. Proteins were transferred onto nitrocellulose membranes using the iBlot 2 Gel Transfer Device. Membranes were then blocked using 5% (wt/vol) nonfat dry milk in tris-buffered saline for 1 hour at room temperature. Blots were subsequently incubated overnight at 4°C with primary antibodies diluted in tris buffered saline containing 0.1% Tween-20 and 4% nonfat dry milk. Blots were then washed with tris buffered saline containing 0.5% Tween-20, and then incubated with secondary antibodies diluted in the same formulation as primary antibodies. Fluorescent signals were then imaged and analyzed using the Li-cor Odyssey DLx Imager and Image Studio software, respectively (LI-COR Biotechnology, Lincoln, NE).

### Pseudovirion and virus-like particle production

HPV16 PsVs were prepared as previously described(61,62). Wild-type HPV16 particles are comprised of capsids formed by L1 and L2 proteins, which encapsidate HPV genomes. Pseudovirions, however, are comprised of L1 and L2 capsid proteins encapsidating reporter plasmids, while virus-like particles are empty capsids made of HPV16 L1 and L2 proteins alone. Briefly, HEK293T cells were co-transfected with codon- optimized HPV16 L1 and L2 p16sheLL plasmid as well as pCIneoGFP reporter plasmid. For bulk PsV preparations, the self-packing p16L1L2 plasmid was utilized (all kind gifts from J. Schiller, Center for Cancer Research, National Institutes of Health, Bethesda, MD). Infectious titer was determined by flow cytometric analysis of fluorescence expression in HEK293T cells 48 h post-treatment with HPV16 PsVs and calculated as IU/mL. Bulk PsV preps were quantified for protein abundance via Coomassie blue staining of diluted PsVs against BSA standards. HPV16 VLPs were produced using a recombinant baculovirus expression system in insect cells as previously described(63).

### Antibodies

Anti-HPV16 L1 antibodies H16.V5 and H16.56E used for immunofluorescence experiments were kind gifts from Neil Christensen (Penn State Cancer Institute, Hershey, PA) and Martin Sapp (Feist-Weiller Cancer Center, Shreveport, LA) respectively(64,65). Anti-WAVE1 (PA5- 78273), anti-WAVE2 (PA5-60975), anti-ITGβ4 (MA5-17104), anti-GAPDH (1D4), Texas Red-X goat anti-rabbit (T6391), Texas Red-X goat anti-mouse (T862), Alexa Fluor 488 goat anti-mouse (A11029), Alexa Fluor 488 goat anti-rabbit (A11034), and Alexa Fluor 680 goat anti-mouse (A21058) antibodies were purchased from ThermoFisher Scientific. Goat anti-mouse IRDye 800CW (925–322) used for Western blotting and imaging was purchased from Li-Cor. Mouse IgG isotype control (ab37355) and rabbit IgG isotype control (ab37415) were purchased from Abcam.

### Post-transcriptional gene silencing

RNAi was conducted by the reverse transfection method. Targets and siRNAs that were used in this study were as follows: Integrin β4 (SI02664102, Qiagen) WASP (S1 - s14835, S2 - s14836 S3 - s14837, ThermoFisher Scientific) WAVE1 (S1 - SI00057946, S2 - SI03022222, S3 - SI03110051, Qiagen) and WAVE2 (S1 - s19802, S2 - s19803, S3 - s19804, ThermoFisher Scientific). For each target gene, three unique, non-overlapping, non-pooled siRNAs (2 μL; 50 μM) were added to individual wells of a 6-well microplate. Silencer Select Negative Control #2 (Ambion) and Allstars Hs Cell Death Positive Control (Qiagen) siRNAs were added to microplates as well to normalize sample wells and assess transfection efficiency, respectively. Lipofectamine RNAiMAX transfection reagent (0.10 μL) (Invitrogen) was added in 1 mL serum-free, antibiotic-free media to microplates containing siRNA. Microplates were incubated for 45 min at room temperature to allow for the sufficient formation of siRNA-to-lipid complexes. 1E5 cells in 1 mL antibiotic-free media containing 20% FBS were added to microplates. The final 2 mL per well containing cells and 50 nM siRNA in antibiotic-free media with 10% FBS was cultured for 48 h at 37°C with 5% CO_2_ and 95% relative humidity before PsV infection assays. Protein knockdown was confirmed at 72, 96, and 120 h post-transfection by Western blotting, which covered the timespan of infection assays.

### Pseudovirus infection assay

Infection is defined in this manuscript as gene transduction and expression of GFP encoded by the reporter plasmid. 2E4 cells (WT, knockout, knockdown, or KO cells overexpressing WAVE1) were seeded in 24-well plates and infected with a 30% tissue culture infective dose (TCID_50_) of PsVs 24 h post-seeding. The percentage of cells expressing the reporter was determined 48 h post-infection via flow cytometry (FC500, Beckman Coulter). TCID_50_ was determined by titrating the multiplicity of infection (MOI) of PsVs to result in approximately 30% infected cells 48 h post-infection.

### Cell surface binding assay

2E5 cells were seeded in 6-well plates and grown overnight. Cells were placed at 4°C for 30 min prior to washing with ice cold PBS supplemented with 1 mM CaCl_2_ as previously described(33). Cells were treated with 10 μg/1E6 cells of HPV16 VLPs in ice cold serum-free media for 1 hour at 4°C to reach binding saturation. Cells were collected on ice via scraping and cell surface VLPs were stained with H16.V5 (1:100) for 30 min at 4°C prior to fixation with 2% paraformaldehyde (PFA). Mean fluorescence intensity (MFI) was used to quantify cell surface binding via flow cytometry.

### Virus internalization assays

2E4 cells were seeded in 24-well microplates and incubated overnight prior to the addition of 2 μg/1E6 cells of HPV16 VLPs conjugated to pHrodo (10:1 dye:HPV L1 ratio) (ThermoFisher Scientific). pHrodo labelled particles were generated using the manufacturer’s protocol and were purified with 2% agarose beads (sized 50-150 μm) (Gold Biotechnology). pHrodo is used as a marker for endocytic trafficking studies because it is a pH-dependent rhodamine dye that is colorless at neutral pH but emits increasing fluorescence as pH decreases. MFI was determined every hour for 0-7 h using the Clariostar plate reader. Microplates were incubated at 37°C with 5% CO_2_ and 95% relative humidity between reads.

### Immunofluorescence microscopy assays

1.5E4 cells were seeded and incubated overnight in 8- well chamber slides with #1.5 polymer coverslip bottoms and ibiTreat surface modification for improved cell attachment (Ibidi). For studies on actin dynamics, cells were treated with CellLight Actin-GFP, BacMam 2.0 at the time of seeding (Invitrogen) or stably transduced with pCMV– LifeAct–TagGFP2 (Ibidi). 24 hours after seeding cells in slides, slides were placed at 4°C for 30 min prior to washing with ice cold PBS supplemented with 1mM CaCl_2_. Cells were then treated with 10 ng/1E6 cells of HPV16 VLPs for 0.5, 1, and 2 h prior to fixation using 4% PFA. For studies examining the relationship between HPV16 and WAVE proteins, cells were then permeabilized with 0.1% Triton X-100 prior to blocking using 1% BSA. For studies of HPV16-stimulated actin protrusions, cells were not permeabilized. HPV16 VLPs were immunostained with H16.5A (1:100). Cells were also stained with the Hoescht 33342 counterstain (1:3000) (ThermoFisher Scientific). For endocytic trafficking colocalization studies, cells were treated with VLPs as above for 0, 2, 4, and 8 hours prior to fixation. Cells were then treated with 0.1% Triton X-100 prior to blocking using 1% BSA.

### Confocal fluorescence microscopy and image analysis

Fluorescence and immunofluorescence associated with cells under different experimental conditions was visualized using a Nikon Eclipse Ti-2 laser scanning confocal microscope equipped with 405, 488, 561, and 640 nm lasers.

Images were analyzed using Imaris software (Oxford Instruments, Abingdon, England). Filopodia were measured manually using the measurement tool. Dorsal surface filopodia were identified as those directly above the nuclear stain and were counted if they protruded >1 µm above the cell surface and within the perimeter of the nuclear stain as viewed from the XY orientation of images. More details on analysis are provided within Fig legends 5-7.

### Statistics

Background from control groups was subtracted in all experiments. All groups were normalized to WT cells or scramble negative control for siRNA experiments for comparison. Statistical analyses were performed using GraphPad Prism 10.0.0 (La Jolla, CA).

## Data availability

The datasets generated during the current study are available from the corresponding author upon reasonable request.

## Acknowledgements

W.M. Kast holds the Walter A. Richter Cancer Research Chair, and this research project was funded by his National Institutes of Health Grant R01 CA074397 and R01 CA074397-S1. Financial contributions through a gift of R.F. Brennan are gratefully acknowledged. The authors would also like to acknowledge the technical contributions of Diane Da Silva, Kim Lühen, Joseph Skeate, and Julia Taylor toward virological studies, and Seth Ruffins for his image analysis expertise.

## Supporting Information

**S1. HPV16 infection in control conditions and western blot for positive control.** (A) Please see Fig 1 for a detailed description of knockdown and infection conditions. Here, we compare HPV16 infection between our scrambled siRNA negative control (NC) with infection in HeLa cells untransfected with siRNA (WT) and cells treated with siRNA against integrin β4, a subunit of a known entry receptor of HPV16 whose knockdown results in ablation of the complete receptor. This served as a positive control in all virological assays involving siRNA knockdown. (B) Western blot protein analysis of integrin β4 (ITGB4) at the time samples were collected to analyze infection.

**S2. Colocalization of HPV16 to endocytic trafficking markers.** Here, we infected WT, W1KO, W2KO, and W1/W2KO HeLa cells with HPV16 VLPs for 8 hours and co-stained for either (A) EEA1, the early endosome marker; (B) VPS25, a marker of multivesicular bodies; (C) Golgin97, a Golgi apparatus marker; and (D) SERCA2, an endoplasmic reticulum marker. Colocalization was determined using the Imaris overlapped volume ratio feature.

**S3. Dorsal surface actin protrusion measurements example.** Here, A WT HeLa cell infected for 0.5 hours with HPV16 PsVs (10 ng/1E6 cells) is depicted. Protrusions were quantified by using the measurement tool in Imaris.

## Notes

### Competing Interest Statement

The authors have declared no competing interest.

