## Supplementary figures and images for "WAVE1 and WAVE2 facilitate human papillomavirus-driven actin polymerization during cellular entry"

### Supplemental Figure 1

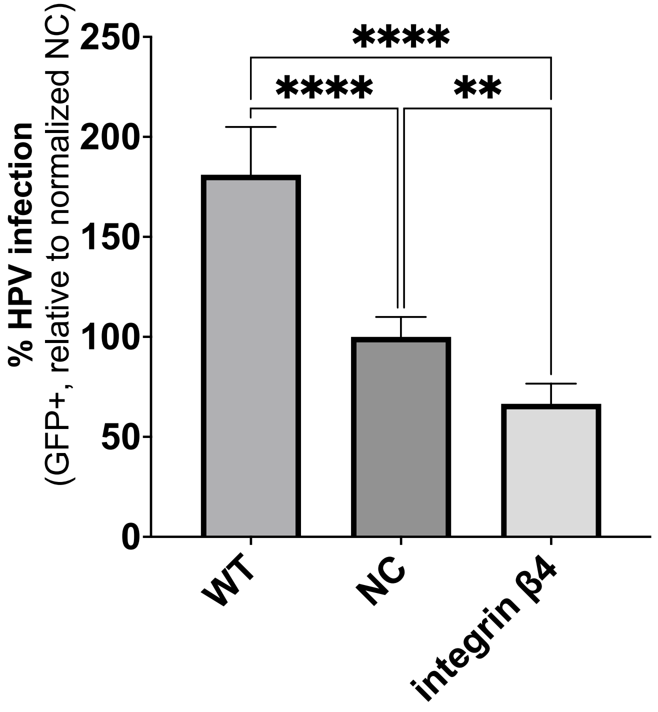

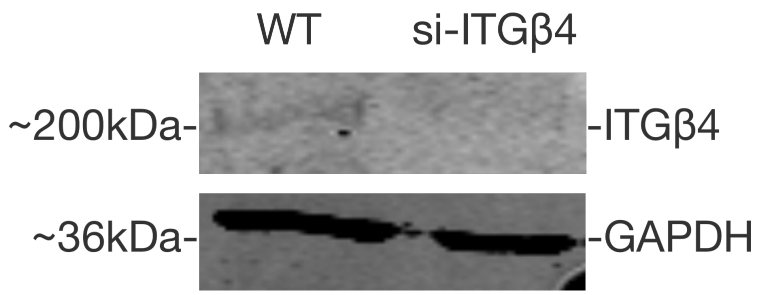

### Supplemental Figure 2

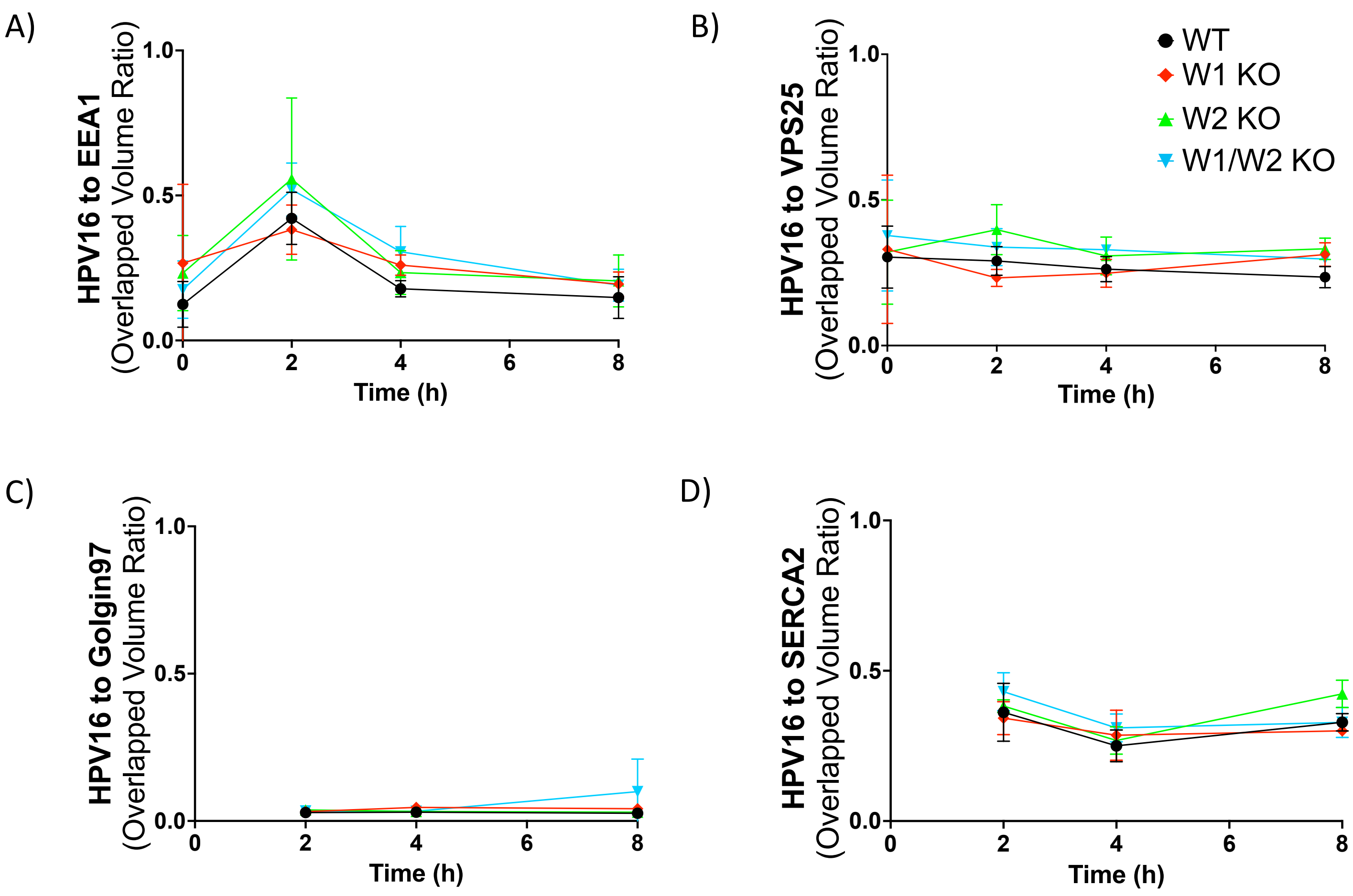

### Supplemental Figure 3

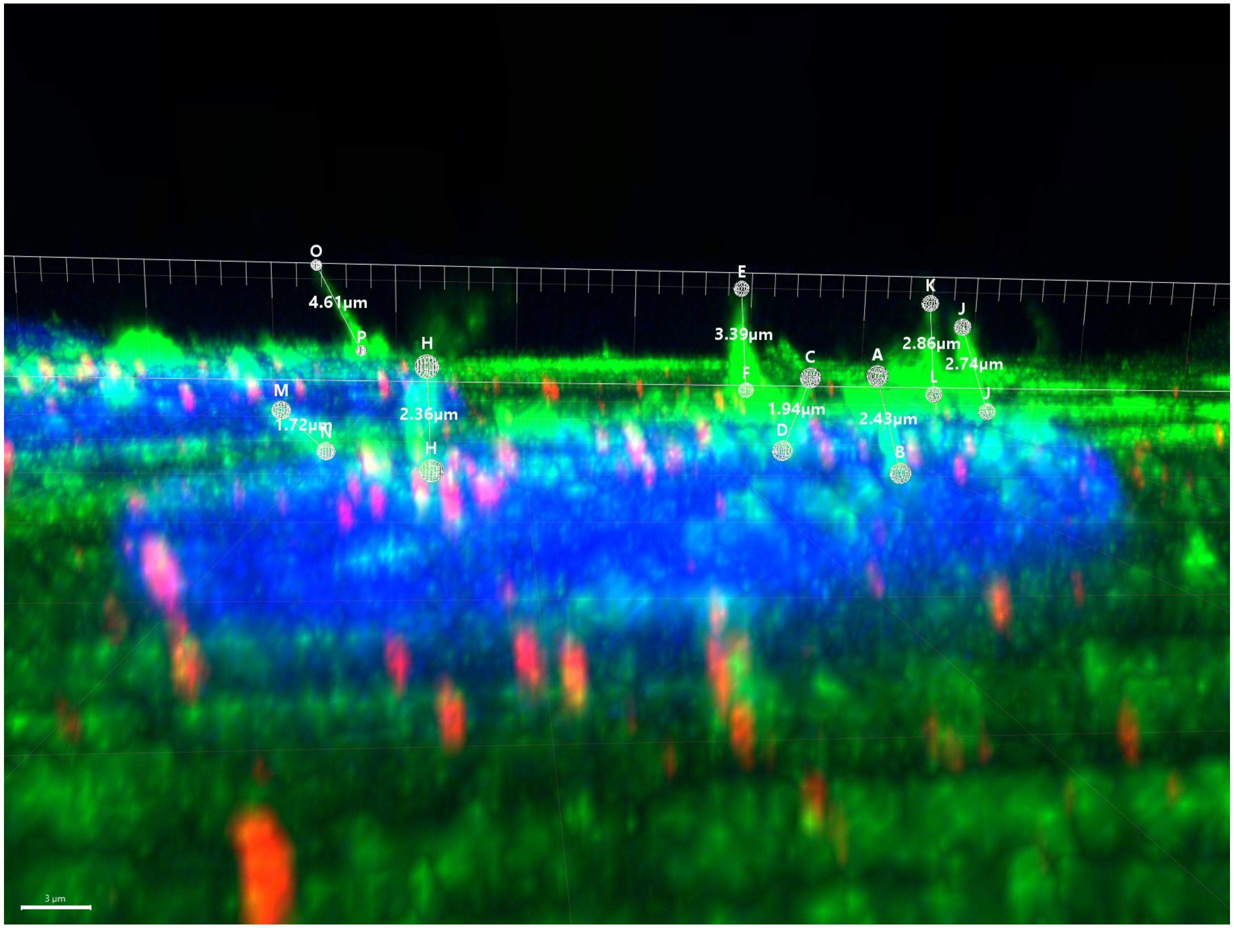
